# Latent gammaherpesvirus exacerbates arthritis and requires age-associated B cells

**DOI:** 10.1101/2021.02.02.429395

**Authors:** Isobel C. Mouat, Zach J. Morse, Iryna Shanina, Kelly L. Brown, Marc S. Horwitz

**Author notes:** Corresponding author: Marc S. Horwitz, Room 3551, Life Sciences Centre, 2350 Health Sciences Mall, University of British Columbia, Vancouver, B. C. Canada V6T 1Z3.

## Abstract

Epstein-Barr virus (EBV) infection is associated with rheumatoid arthritis (RA) in adults, though the nature of the relationship remains unknown. Herein, we examine the contribution of viral infection to the severity of arthritis in mice. We provide the first evidence that latent gammaherpesvirus infection enhances clinical arthritis, modeling EBV’s role in RA. Mice latently infected with a murine analog of EBV, gammaherpesvirus 68 (γHV68), develop more severe collagen-induced arthritis and a Th1-skewed immune profile reminiscent of human disease. We demonstrate that disease enhancement requires viral latency and is not due to active virus stimulation of the immune response. Age-associated B cells (ABCs) are associated with several human autoimmune diseases, including arthritis, though their contribution to disease is not well understood. Using ABC knockout mice, we provide the first evidence that ABCs are mechanistically required for viral enhancement of disease, thereby establishing that latent gammaherpesvirus infection stimulates ABCs to provoke arthritis.

**Conflict of interest statement:** The authors have declared that no conflict of interest exists.

## Introduction

Rheumatoid arthritis (RA) is one of the most common autoimmune diseases in adults, though the etiology and pathophysiology are not fully understood(1,2). RA, as well as other autoimmune diseases including multiple sclerosis and systemic lupus erythematosus, is associated with Epstein-Barr virus (EBV) infection(3–5). The circulating EBV load is higher in individuals with RA than otherwise healthy adults(3) and RA patients have increased levels of antibodies specific to multiple EBV-encoded proteins(6–10). Further, RA patients have increased EBV-specific CD8^+^ T cells(11) yet these cells have a reduced ability to kill EBV-infected B cells when compared to the same subset of EBV-specific CD8^+^ T cells from healthy controls(12). However, the precise role of EBV in RA pathogenesis remains unknown. EBV infection typically takes place during childhood or adolescence, while RA generally becomes symptomatic during middle age, indicating that the latent EBV infection likely modulates the immune system over time in a manner that contributes to the development of RA(1,13–15).

Evidence from in vivo models are scarce and previous studies have focused primarily on the direct relationship between EBV infection and damage to the joint capsule, with little attention given to systemic effects of EBV infection on immune modulation preceding and continuing throughout disease. Mice with humanized immune systems, namely NOD/Shi-*scid*/IL-2Rγ^null^ mice reconstituted with CD34^+^ hematopoietic stem cells, that were infected with EBV went on to spontaneously develop erosive arthritis, suggesting a causative role of EBV in arthritis development(16). Related, a serum transfer-induced arthritis model was used to demonstrate that Ly6C^high^ monocytes play a role in transporting murine gammaherpesvirus 68 (γHV68), an EBV homolog, to the synovium(17). Our group has previously shown that latent γHV68 infection enhances experimental autoimmune encephalomyelitis (EAE) and leads to a disease that more closely resembles multiple sclerosis (MS)(18). Critically, this enhancement was specific to γHV68; other viruses, including lymphocytic choriomeningitis virus (LCMV) and murine cytomegalovirus (MCMV), did not lead to enhancement of EAE. Additionally, enhancement took place without changes to autoantibody levels. An in vivo model that recapitulates the temporal and systemic immunological aspects of the relationship between EBV and RA is critical.

To examine the relationship between EBV and RA, we have adapted in vivo models of both. γHV68 is a natural pathogen that is aγHV68 well-established and widely-used murine model of EBV infection that shares an array of characteristics with human EBV infection, including latent persistence in B cells, a potent CD8 T cell response, and immune evasion tactics(19,20). Type II collagen-induced arthritis (CIA) is a commonly used model of RA wherein mice are injected with type II collagen emulsified in adjuvant. Here, we chose to use C57Bl/6 mice due to the extensive past characterization of γHV68 infection in C57Bl/6 mice and the numerous knockout strains available on this background. Multiple strains of mice are susceptible to CIA, including C57Bl/6 mice that, despite displaying a less severe disease course than other strains, generate a robust T cell response. In C57Bl/6 mice, CIA follows a chronic disease course with a sustained T cell response, presence of anti-collagen IgG, and infiltration of inflammatory lymphocytes into the joint capsule(21). Here we show that C7Bl/6 mice latently infected with γHV68 and induced for CIA develop a more severe clinical course and altered immunological profile, with an expansion of CD8^+^ T cells and Th1 skewing. We have utilized γHV68 infection and CIA induction to investigate the mechanism(s) by which EBV contributes to RA, in particular the contribution of age-associated B cells.

The role of B cells the contribution of EBV to RA is intriguing because B cells contribute pathogenically to RA, and EBV infects B cells and alters the B cell profile(22,23). Age-associated B cells (ABCs) are a subset of B cells that are of particular interest as they have been implicated in both autoimmunity and viral infection. When compared to healthy adults, the relative proportion and/or absolute circulating counts of ABCs are elevated in RA patients, a subset of individuals with MS, individuals with systemic lupus erythematosus (SLE), and a subset of people with common variable immune deficiency that display autoimmune complications’(24–31). ABCs are required for disease development in mouse models of SLE(32). Also, ABCs are increased during viral infections in mice and/or humans including lymphocytic choriomeningitis virus, γHV68, vaccinia, HIV, hepatitis C virus, and influenza. ABCs display an array of functional capacities, including the secretion of anti-viral or autoantibodies, initiation of germinal centres, antigen presentation to T cells, and secretion of cytokines(33–36). It is yet to be examined whether ABCs play a role in viral contribution to autoimmunity. We find that ABC knockout (KO) mice are unable to develop the γHV68-exacerbation of CIA, and therefore act as a mediator between viral infection and autoimmunity.

## Results

### Latent γHV68 infection exacerbates the clinical course of CIA

The development of RA often occurs years after initial infection with EBV when the virus is latent. To mimic this temporal relationship, we infected mice with γHV68, waited five weeks for the lytic infection to clear and the virus to establish latency, and induced CIA. Clearance of the acute virus and establishment of latency has previously been shown by plaque assay on spleens collected 35 days post-infection(18,37). Following CIA induction, mice were assessed three times per week for redness and swelling in the hind two paws (**Figure 1 – figure supplement 1A**), which informed a clinical score (5-point scale from 0-4) for each mouse. We observe that CIA in latent γHV68-infected mice (herein referred to as γHV68-CIA mice) has a more severe clinical course than uninfected mice (herein referred to as CIA mice), as evidenced by consistently higher clinical scores at disease onset and over the following 8 weeks (**Figure 1A**; fold change-1.4 at day 56 post-induction). γHV68-CIA mice also develop onset of disease symptoms an average of seven days earlier than CIA mice (**Figure 1C**). In agreement with other research groups, male and female mice displayed similar clinical scores during CIA and we also do not observe a sex difference in γHV68-CIA mice (**Figure 1 – figure supplement 1B**). As expected, latent γHV68 infected mice (without CIA) did not display any signs of disease (**Figure 1A-B**). Titers of anti-type II collagen autoantibodies (total IgG, IgG1, and IgG2c) were elevated in sera from mice with CIA compared to naïve mice without CIA yet were similar in mice with CIA regardless of infection (**Figure 1 – figure supplement 1C-E**), in line with our previous finding that latent γHV68 infection enhances EAE without influencing autoantibody levels(18). These data demonstrate that latent γHV68 infection leads to earlier onset and more severe CIA, though the exacerbation is not due to higher titers of autoantibodies against type II collagen or change in abundance of particular immunoglobulin isotypes.

**Figure 1:**
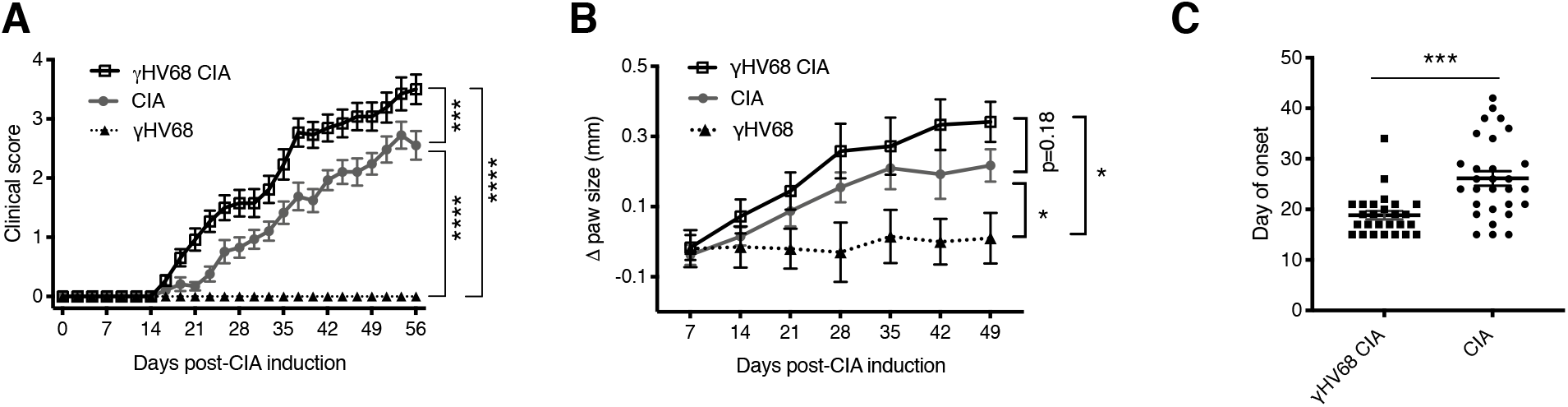
Progression of CIA in latent γHV68 infected and control uninfected mice. (**A**) Clinical score (y-axis) of CIA measured three times weekly for 8 weeks (x-axis; days) post CIA induction in mice without (CIA, filled circles) and with latent γHV68 infection (γHV68-CIA, open squares), and starting at day 35 post-infection in mice infected with latent γHV68 infection but not induced for CIA (γHV68, filled triangles). (**B**) Change (Δ, y-axis) in thickness of hind paws measured with calipers once per week and averaged for each mouse. Began measuring γHV68-CIA and CIA the day of CIA induction, and γHV68 at day 35 post-infection. (**C**) Day (y-axis) of CIA onset (considered two consecutive scoring days of a score of at least 1) in mice (x-axis) without (CIA) and with latent γHV68 infection (γHV68-CIA). Each data point represents an individual mouse. (**A-C**) Data presented as mean ± SEM. Statistical significance determined by (**A, B**) two-way ANOVA with Geisser-Greenhouse’s correction, (**C**) t-test. *** p<0.001, ** p<0.01, * p<0.05. (**A**) n=10-29 mice per group, 4 experiments; (**B**) n=8-20 mice per group, 3 experiments; (**C**) n=26-29 mice per group, 4 experiments.

### The profile of immune cells infiltrating the synovium is altered in γHV68-CIA

To assess the types and relative proportions of immune cells infiltrating the joint synovium, synovial fluid cells were collected on day 56 post-CIA induction. Synovial cells were collected from the knee and ankle joints by flushing each joint with PBS and subsequently analyzing isolated cells by flow cytometry. Synovial cells were not collected from naïve or γHV68-infected mice without CIA because we would not expect there to be sufficient infiltration of immune cells for analysis. The number of CD8^+^ T cells infiltrating the synovium during γHV68-CIA is increased compared to CIA (3.6-fold change), while there is no significant difference in the number of CD4^+^ T cells (**Figure 2A**). Additionally, the CD8^+^ and CD4^+^ T cells in γHV68-CIA synovium display a significant increase in Tbet expression compared to those in CIA (**Figure 2B**), indicating Th1 skewing.

**Figure 2:**
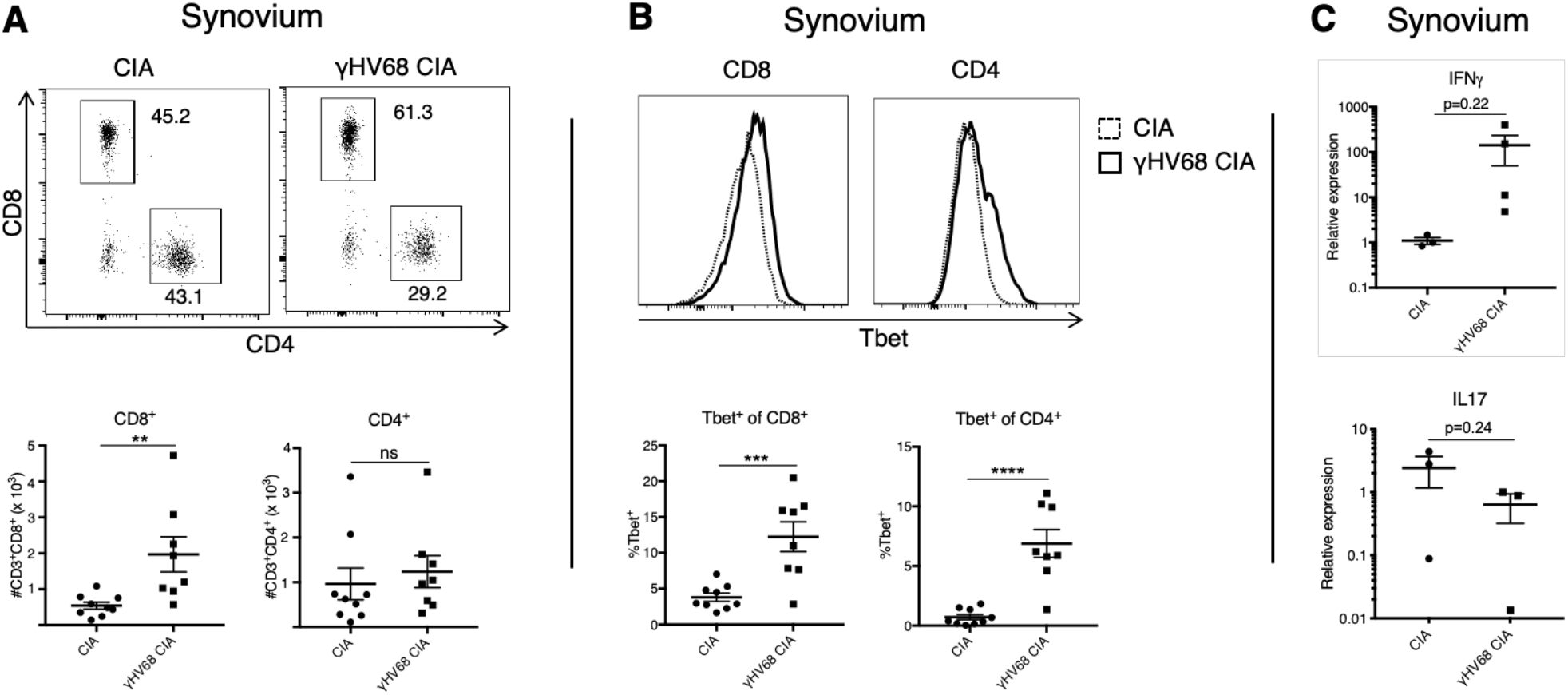
Analysis of immune infiltration to synovium between γHV68-CIA and control CIA mice at day 56 post-CIA induction. (**A**) Representative flow cytometry plots of synovial fluid (SF) CD8^+^ and CD4^+^ T cells. Previously gated on lymphocytes, singlets, live cells, and CD45^+^CD3^+^ cells. Total numbers (y-axis) of CD8^+^ and CD4^+^ T cells in uninfected mice with CIA (filled circles) and γHV68-CIA mice (filled squares). (**B**) Representative flow cytometry plots of Tbet expression (x-axis) by CD8^+^ or CD4^+^ T cells in CIA mice (dotted line) and γHV68-CIA mice (solid line). Samples previously gated on lymphocytes, singlets, live cells, and CD45^+^CD3^+^ cells. Percent of CD8^+^ and CD4^+^ T cells positive for Tbet (y-axis, gated on a full minus-one control) in uninfected mice with CIA (filled circles) and γHV68-CIA mice (filled squares). (**C**) RNA extracted from synovial fluid cells, RT-qPCR performed for IFNγ and IL17, and relative expression plotted for uninfected mice with CIA (filled circles) and γHV68-CIA mice (filled squares). (**A-B**) Flow plots are concatenated samples from all CIA or γHV68-CIA samples from an individual experiment, n=8-9 mice per group; (**C**) n=3-4 mice per group; (**A-C**) One experiment, data presented as mean ± SEM, analyzed by t-test, ****p<0.0001, *** p<0.001, ** p<0.01, * p<0.05.

As further evidence that infiltrated T cells are immunologically active, we used RT-qPCR to evaluate the expression of key T cell derived cytokines IFNγ and IL17. The relative expression of IFNγ in synovium cells of γHV68-CIA mice compared to CIA-mice was increased (fold chage-129), while the relative expression of IL-17A is trending down in infected mice (**Figure 2C;** fold change-3.8). Together, these results indicate that IFNγ-producing T cells are preferentially infiltrating the synovium in our model of γHV68-CIA, which is consistent with what is observed in the synovium of RA patients(38). Our data also demonstrate a skewing towards cytotoxic CD8^+^ T cells in mice latently infected with γHV68 prior to CIA.

### Latent γHV68 infection skews the T cell response towards a pathogenic profile during CIA

To examine how latent γHV68 might contribute to CIA, we specifically examined the systemic T cell profile. It is known that latent γHV68 infection expands cytotoxic T cells and reduces Tregs(39). Both cell types play a role in CIA with cytotoxic T cells being crucial mediators of CIA while Tregs play a protective role(40,41). We examined T cells in the spleen and inguinal lymph nodes (ILNs), a draining lymph node in which we observe a significant increase in overall abundance of immune cells during CIA (data not shown). γHV68-CIA mice display a decrease in relative proportion of FoxP3^+^ Tregs and an increase in relative proportion of CD8^+^ T cells in the spleen compared to control CIA mice (**Figure 3A-B, Figure 3 – figure supplement 1A-B**). This is similar to what is observed in people with RA, as activated CD8^+^ T cells are increased and Tregs are decreased in the circulation of RA patients compared to otherwise healthy people(42,43). In the ILNs of γHV68-CIA mice, we observe a trend of decreased relative proportion of regulatory T cells and also decreased CD8^+^ and CD4^+^ T cells relative proportions, indicating potential T cell egress from the ILNs during disease (**Figure 3A-B, Figure 3 – figure supplement 1D-F**). We also observe a significant increase in relative proportion of CD11c^+^CD8^+^ DCs in γHV68-CIA mice compared to CIA (**Figure 3 – figure supplement 1D-F**). This data shows that the T cell profile of γHV68-CIA mice is skewed pathogenically, with decreased Tregs and increased cytotoxic T cells.

**Figure 3:**
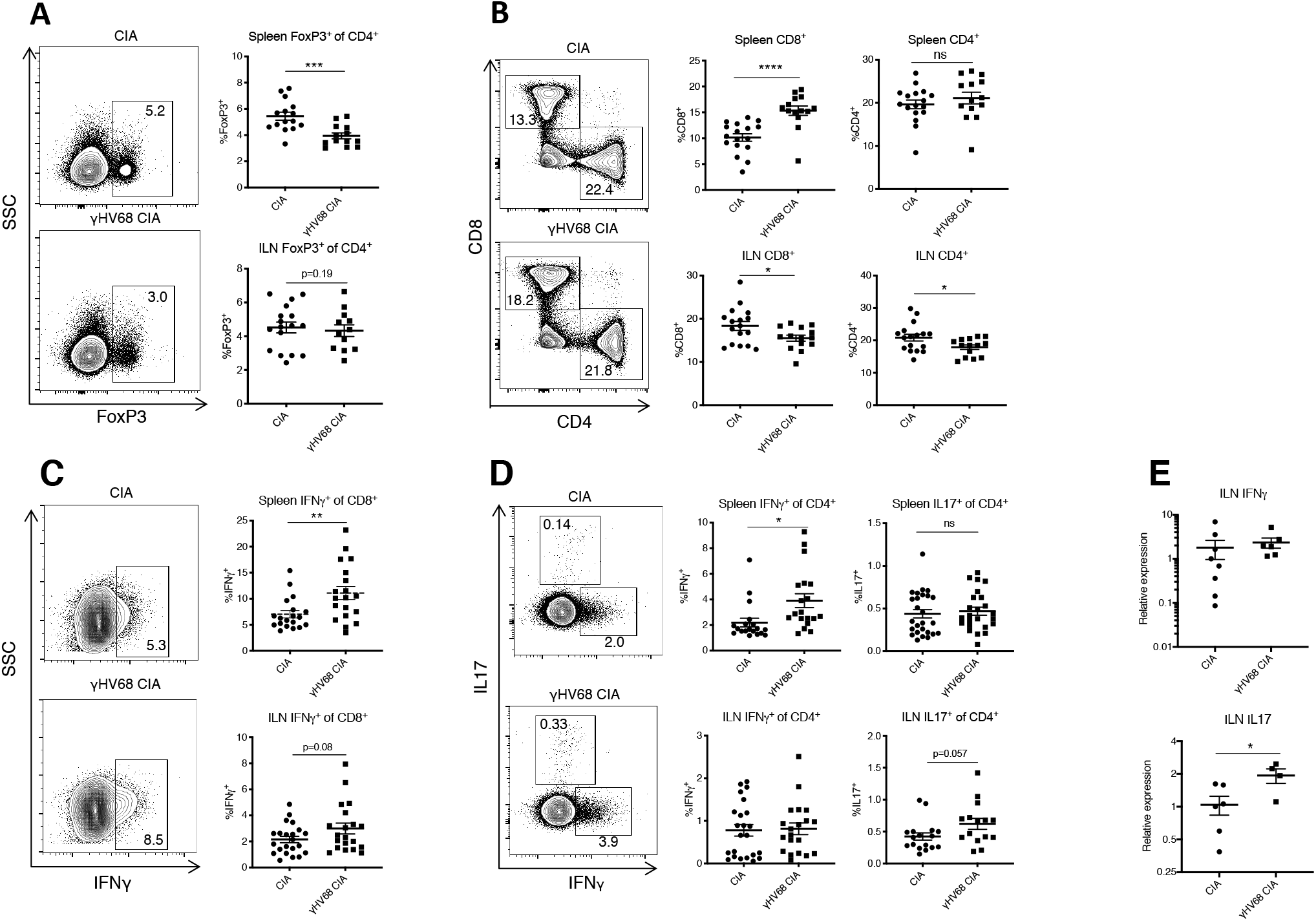
Flow cytometry analysis of spleen and ILN T cells at day 56 post-induction of γHV68-CIA and control CIA mice. (**A-D**) Representative flow cytometry plots of spleen samples previously gated on lymphocytes, live cells, singlets, CD45^+^CD3^+^ cells, and (**A, D**) CD4^+^ cells, (**C**) CD8^+^ cells, of uninfected mice with CIA (upper plot) and γHV68-CIA mice (lower plot). (**A, C**) Side-scatter (SSC) plotted on the y-axis. (**A-D**) Percent of immune subsets (y-axis) in the spleens of uninfected mice with CIA (filled circles) and γHV68-CIA mice (filled squares). **(A)** %FoxP3^+^ of CD4^+^; (**B**) %CD3^+^CD8^+^ and %CD3^+^CD4^+^ of CD45^+^; (**C**) %IFNγ^+^ of CD8^+^; (**D**) IFNγ^+^ or IL17^+^ of CD4^+^; (**E**) RNA extracted from ILN cells, RT-qPCR performed for IFNγ and IL17, and relative expression plotted for uninfected mice with CIA (filled circles) and γHV68-CIA mice (filled squares). (**A**) n=12-17 mice per group, 3 experiments; (**B**) n=14-17 mice per group, 3 experiments; (**C, D**) n=19-22 mice per group, 3 experiments; (**E**) n=4-8 mice per group, 1 experiment. (**A-E**) Each data point represents an individual mouse. Data presented as mean ± SEM, analyzed by t-test, ****p<0.0001, *** p<0.001, ** p<0.01, * p<0.05.

### T cell polarization is modulated in γHV68-CIA mice

Although IL17 has been highly studied due to its predominance in animal models of arthritis, both IL17 and IFNγ are involved in RA(38,43–45). As expected from our previous work with γHV68-EAE, we find that in γHV68-CIA greater numbers of splenic CD8^+^ and CD4^+^ T cells express IFNγ compared to CIA alone (**Figure 3C-D**). There is a maintenance of Th17 cells in the spleen, with a similar proportion of CD4^+^ T cells expressing IL17 in CIA and γHV68-CIA (**Figure 3D**). In the ILNs, we observe a significant increase in IL17 by RT-qPCR (**Figure 3E**) and a corresponding trend towards more IL17-expressing CD4^+^ T cells. We propose that the combined Th1 and Th17 profile observed in γHV68-CIA is more reminiscent of what is observed in people with RA than in CIA without γHV68 infection.

### Latency is required for the clinical and immunological γHV68-exacerbation of CIA

To examine the requirement of γHV68 latency, as opposed to residual effects from acute infection, for exacerbating CIA, we used a recombinant γHV68 strain that does not develop latency, ACRTA-γHV68. In ACRTA-γHV68 the genes responsible for latency are deleted and a lytic gene, RTA, is constitutively expressed, resulting in clearance of the virus following acute infection(46). We find that ACRTA-γHV68 infected mice do not develop the CIA clinical enhancement that we observe in latently γHV68-infected mice, with the clinical course and day of onset resembling that of uninfected CIA mice (**Figure 4A-B**).

**Figure 4:**
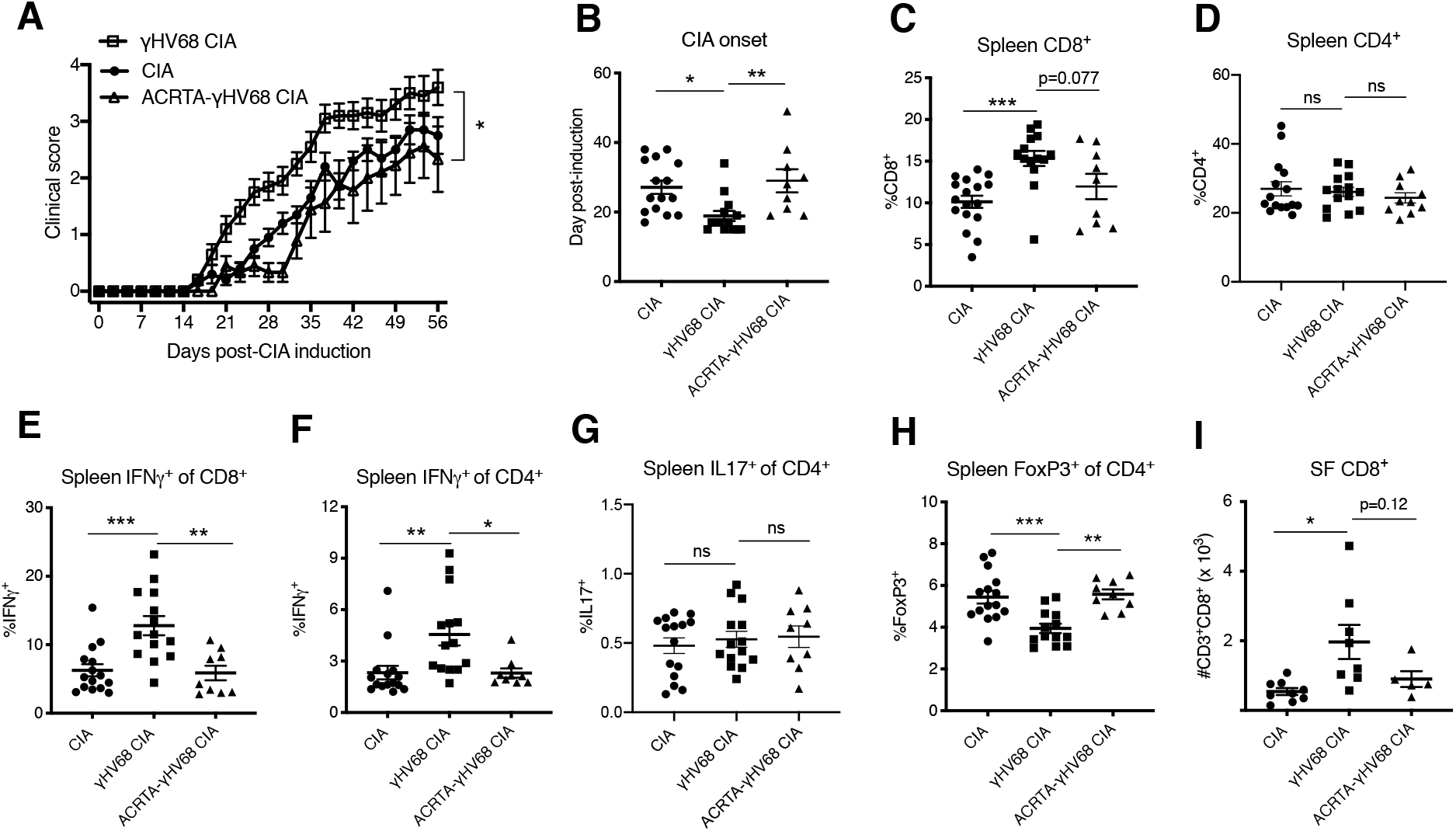
Disease progression and immune profile of latency-free ACRTA-γHV68 CIA mice compared to γHV68-CIA and CIA mice. CIA was induced in C57BL/6 mice after 5 weeks of mock, γHV68, or ACRTA-γHV68 infection, and mice scored for clinical disease until 56 days-post CIA induction. At 56 days post CIA-induction, spleens and synovial fluid collected and processed for flow cytometry. A proportion of the CIA and γHV68-CIA data is repeated from Figure 2. (**A**) Clinical scores (y-axis) of uninfected mice with CIA (filled circles), γHV68-CIA mice (open squares), and ACRTA-γHV68 CIA mice (open triangles); (**B-I**) Comparison of uninfected mice with CIA (filled circles), γHV68-CIA mice (filled squares), and ACRTA-γHV68 CIA mice (filled triangles). (**B**) Day (y-axis) of CIA onset, considered two consecutive scoring days of a score of at least 1, in mice (x-axis) without (CIA) and with latent γHV68 infection (γHV68-CIA) or ACRTA-γHV68 infection (ACRTA-γHV68 CIA). (**C-I**) Immune cell subsets determined by flow cytometry, previously gates on lymphocytes, singlets, live cells, and CD45^+^ cells; (**C**) %CD3^+^CD8^+^ of CD45^+^ cells in the spleen; (**D**) %CD3^+^CD4^+^ of CD45^+^ cells in the spleen; (**E**) %IFNγ^+^ of CD8^+^ cells in the spleen; (**F**) %IFNγ^+^ of CD4^+^ cells in the spleen; (**G**) IL17^+^ of CD4^+^ cells in the spleen; (**H**) %FoxP3^+^ of CD4^+^ cells in the spleen; (**I**) Number of CD3^+^CD8^+^ cells in synovial fluid (SF) determined by flow cytometry. (**A-H**) n=9-15 mice per group, 2 experiments, (**I**) n=5-9 mice per group, 1 experiment. (**A-G**) Each data point represents an individual mouse. Data presented as mean ± SEM. Analyzed by (**A**) two-way ANOVA with Geisser-Greenhouse’s correction, (**B-I**) One-way ANOVA, *** p<0.001, ** p<0.01, * p<0.05, ns = not significant.

Furthermore, the immunological changes observed in γHV68-CIA mice, when compared to CIA mice, are absent in ACRTA-γHV68 CIA mice. The increase in relative proportion of CD8^+^ T cells in the spleen is less pronounced in ACRTA-γHV68 CIA compared to γHV68-CIA, while there is no change in relative proportion of CD4^+^ T cells (**Figure 4A-B**). In ACRTA-γHV68 CIA mice there is abolishment of the γHV68-induced upregulation of IFNγ in CD8^+^ and CD4^+^ T cells, and no change in IL17 expression by CD4^+^ T cells (**Figure 4E-G**). The decrease in relative proportion of splenic Tregs and CD8^+^ infiltration into the synovial fluid observed in γHV68 CIA mice is not present in ACRTA-γHV68 CIA mice (**Figure 4H-I**). Together, this data shows that ACRTA-γHV68 CIA mice display a similar clinical and immunological profile to uninfected CIA mice. This demonstrates that the enhancement is not due to innate immune stimulation during the acute infection, but, rather, the latency phase of γHV68 infection is critical for the clinical and immunological exacerbation of CIA. The requirement of γHV68 latency mirrors the RA patient clinical course, wherein patients are infected with EBV years before the onset of disease.

### Age-associated B cells are increased and display an inflammatory phenotype in γHV68-CIA

As the number of ABCs are expanded in the contexts of both viral infection and autoimmunity, including RA(25,28,29,33,35), we investigated the role of ABCs facilitating in viral enhancement of CIA. We began by examining the proportion and phenotype of ABCs in uninfected CIA mice and CIA mice previously infected with latent γHV68 (γHV68-CIA) by flow cytometry (**Figure 5 – figure supplement 1A**). We find that CIA induction increases the proportion and total number of ABCs (CD19^+^CD11c^+^Tbet^+^) in the spleen, and γHV68-CIA mice have further increased proportions of ABCs in the spleen compared to CIA (**Figure 5A-B**). The proportion of ABCs in the ILNs is not a significantly different between γHV68-CIA and CIA mice (**Figure 5 – figure supplement 1C**). The number of ABCs is substantially lower in the ILNs than in the spleen, concurring with other studies that find ABCs primarily reside in the spleen(36). During CIA and γHV68-CIA, infecting mice at 6-8 weeks of age, we do not observe differences in the proportions of ABCs between male and female mice (**Figure 5 – figure supplement 1B**).

**Figure 5:**
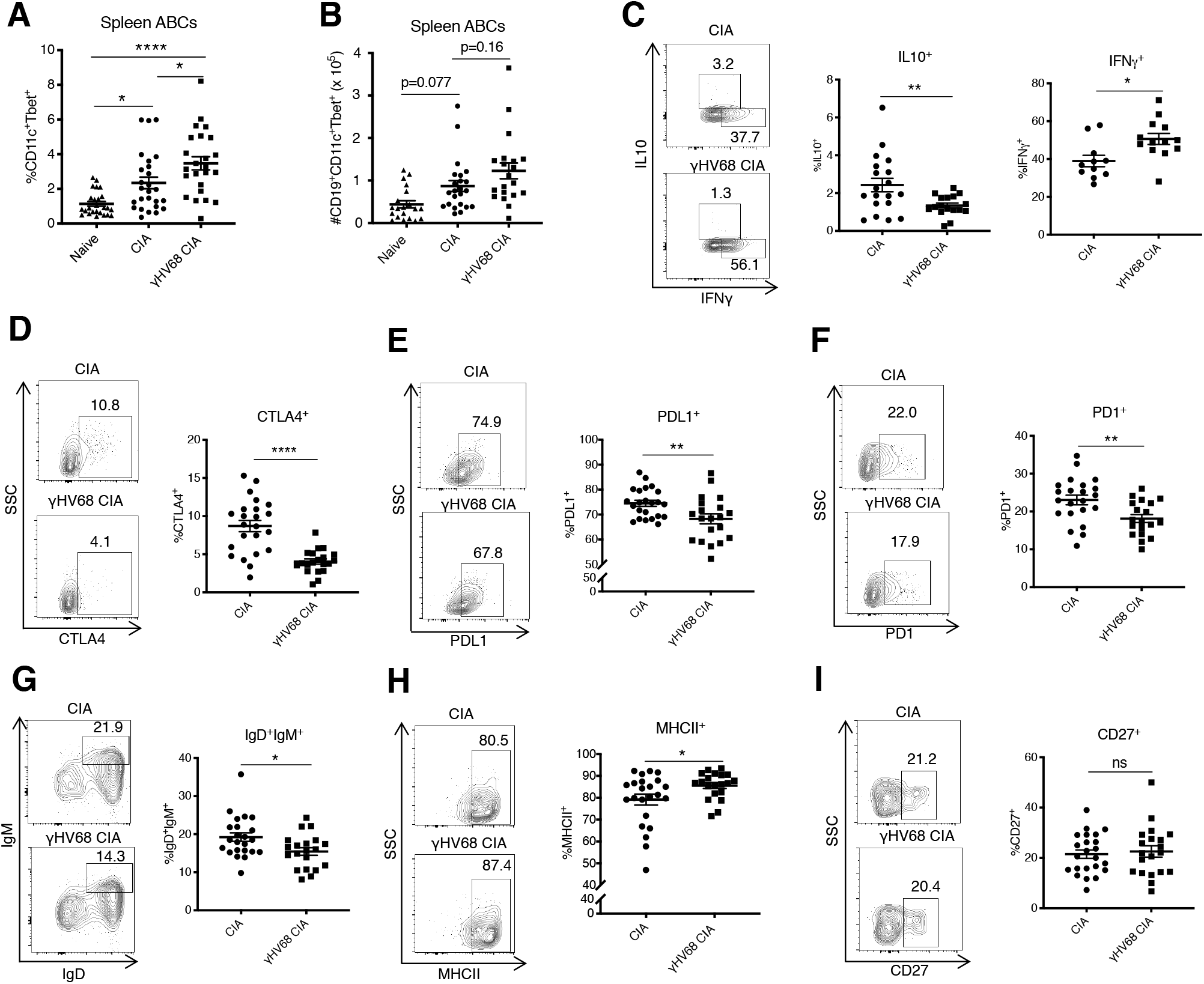
Analysis of ABC amount and phenotype by flow cytometry at 56 days post-CIA induction. (**A**) Percentage of ABCs (CD11c^+^Tbet^+^) of mature B cells (CD19^+^IgD^−^) in the spleen and (B) total numbers of ABCs in the spleen of naïve mice (filled triangle), uninfected mice with CIA (filled circles) and γHV68-CIA mice (filled squares). (**C-I**) Phenotype of ABCs analyzed by flow cytometry. Previously gated on splenic CD19^+^CD11c^+^Tbet^+^ ABCs. Flow plots are representative samples, SSC = side scatter. Proportion of ABCs positive for (**C**) IL10 and IFNγ, (**D**) CTLA4, (**E**) PDL1, (**F**) PD1, (**G**) IgD^+^IgM^+^, (**H**) MHCII, (**I**) CD27. (**A**) n=24-26 mice per group, 3 experiments, (**B**) n=20-23 mice per group, 3 experiments, (**C**) n=16-19 mice per group, 2 experiments, (**D-I**) n=20-23 mice per group, 2 experiments. (**A-I**) Each data point represents an individual mouse. Data presented as mean ± SEM. (**A-B**) Analyzed by one-way ANOVA, (**C-I**) Analyzed by t-test, **** p<0.0001, *** p<0.001, ** p<0.01, * p<0.05.

We next examined the phenotypic characteristics and find that ABCs in the spleen are phenotypically distinct in γHV68-CIA compared to CIA. First, fewer ABCs in the spleens of γHV68-CIA mice express IL10, while an increased proportion express IFNγ (**Figure 5C**), indicating that they are skewed towards a pathogenic Th1 phenotype. Further, fewer splenic ABCs in γHV68-CIA mice express inhibitory receptors CTLA4, PDL1, and PD1 (**Figure 5D-F**). This indicates that ABCs in CIA mice play a more regulatory role than those in γHV68-CIA mice. Additionally, the ABCs in γHV68-CIA mice display a more mature phenotype, with fewer IgD^+^IgM^+^ naïve B cells and increased MHCII expression, though the expression of memory marker CD27 is unchanged (**Figure 5G-I**). These results indicate that ABCs in γHV68-CIA mice are more mature and may have increased antigen presentation capacities but are not primarily a memory subset. There are no differences in the expression of CD20, TNFα, CD95 (Fas), nor IDO expression (**Figure 5 – figure supplement 1D-G**). Collectively these results indicate that ABCs in γHV68-CIA mice display a more pathogenic phenotype than those in CIA, with decreased expression of regulatory cytokine IL10 and inhibitory markers, and increased expression of IFNγ.

### Age-associated B cells are required for γHV68-exacerbation of CIA

To determine whether ABCs are a subset mediating the viral enhancement of CIA we utilized ABC knock-out mice that harbor a B cell specific Tbet deletion. The clinical course and immune profile of CIA and γHV68-CIA mice was compared in littermate controls of Tbet^fl/fl^ × CD19Cre^+/−^ (KO) and Tbet^fl/fl^ × CD19Cre^−/−^ (Ctrl) mice (**Figure 6A**). We observe that the clinical course is unchanged in CIA between Ctrl and KO mice, indicating that ABCs are not contributing to the disease course in CIA (**Figure 6B**). Alternatively, when induced with CIA, γHV68-infected KO mice do not display the γHV68-exacerbated clinical course compared to γHV68-CIA Ctrl mice (**Figure 6C**), indicating that ABCs are a pathogenic subset in γHV680-CIA. Without ABCs, γHV68-CIA mice do not display clinical exacerbation, but rather appear similar to uninfected CIA mice in terms of disease severity and day of onset (**Figure 6D-E**). We observe that the ablation of ABCs results in a decreased relative proportion of splenic CD8^+^ T cells in γHV68-CIA, though there is no difference in relative proportion of CD4^+^ T cells (**Figure 6F-G**). Relative proportion of Tregs are slightly increased in KO mice in both CIA and γHV68-CIA (**Figure 6H**). Expression of IL17 or IFNγ by CD4^+^ or CD8^+^ T cells is unchanged between Ctrl and KO groups (data not shown). These results indicate that ABCs are important for coordinating the pathogenic CD8^+^ T cell response and are a critical pathogenic population in γHV68-CIA.

**Figure 6:**
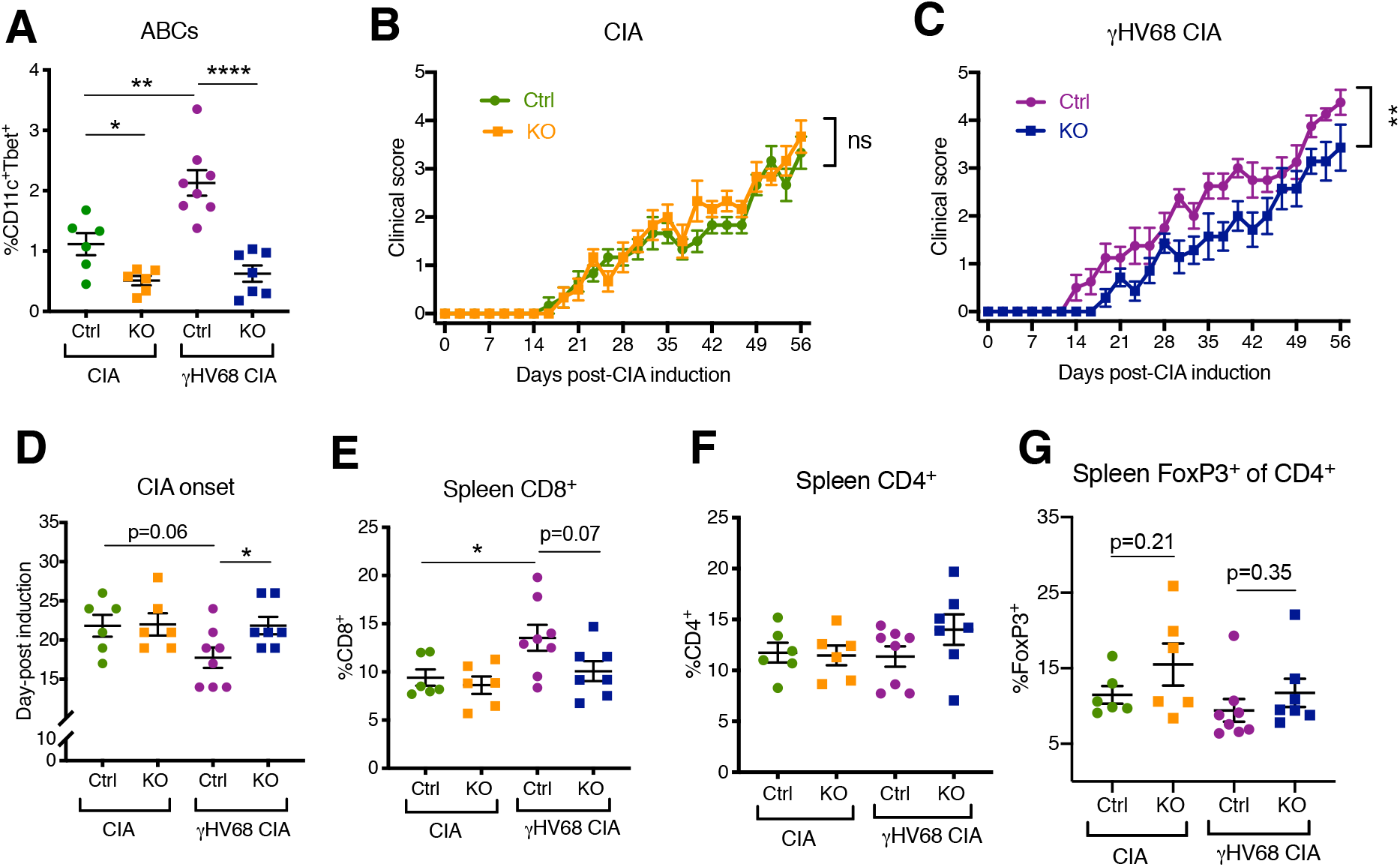
Disease progression and flow cytometric analysis of Tbet^fl/fl^ × CD19Cre^+/−^ mice (KO) compared Tbet^fl/fl^ × CD19Cre^−/−^ mice (Ctrl) that have been infected with γHV68 or mock-infected and induced for CIA. **(A)** Proportion of ABCs (CD11c^+^Tbet^+^) of mature B cells (CD19^+^IgD^−^) in the spleen as determined by flow cytometry of CIA flox-only control (Ctrl, green circles), CIA KO (KO, orange squares), γHV68-CIA flox-only control (Ctrl, purple circles) and γHV68-CIA KO (KO, blue squares) mice. (**B**) Clinical score (y-axis) of CIA measured three times weekly for 8 weeks (x-axis; days) post CIA induction in Ctrl (green circles) and KO (orange circles) mice that are uninfected with CIA. (**C**) Clinical score (y-axis) of CIA measured three times weekly for 8 weeks (x-axis; days) post CIA induction in Ctrl (purple circles) and KO (blue circles) γHV68-infected CIA mice. (**D**) Day (y-axis) of CIA onset, considered two consecutive scoring days of a score of at least 1, in Ctrl and KO CIA and γHV68-CIA mice (x-axis). (**E-G**) T cells examined in the spleen at day 56 post-induction by flow cytometry. Previously gated on lymphocytes, live cells, singlets, and CD45^+^CD3^+^ cells. (**E**) %CD3^+^CD8^+^ of CD45^+^, (**F**) %CD3^+^CD4^+^ of CD45^+^, (**G**) %FoxP3^+^ of CD4^+^ cells in the spleen. (**A-G**) n=6-8 mice per group, 2 experiments. Data presented as mean ± SEM. Analyzed by (**B-D**) two-way ANOVA, (**A, D-G**) t-test, ****p<0.0001, *** p<0.001, ** p<0.01, * p<0.05.

## Discussion

In this report, we demonstrate that latent γHV68 exacerbates CIA clinically and immunologically, and Tbet^+^ B cells, known as age-associated B cells, are critical for this exacerbation. Investigation of the mechanism by which EBV contributes to RA has previously been challenging due to the lack of a murine model to examine the systemic immune modulation caused by latent gammaherpesvirus infection and resulting influence on arthritis. Here, we show that infecting mice with latent γHV68 prior to CIA induction results in an immune course more similar to that of RA patients than CIA alone, and is a suitable model for examining the contribution of EBV to RA. Elucidating how EBV infection contributes to the development of RA is critical to understanding the underlying pathophysiology of the disease.

As EBV is associated with several autoimmune diseases, it is important to examine whether there are conserved mechanisms of contribution. The overlap in etiology and pathophysiology between these autoimmune diseases may help to explain the cross-efficacy of immunotherapies between MS and RA, including B cell depletion therapies. Our lab has previously demonstrated that latent γHV68 infection enhances EAE, a common model of MS, clinically and immunologically(18). In both the γHV68-CIA and γHV68-EAE models we observe an increase in CD8^+^ T cells at the site of disease and increased expression of IFNγ by cytotoxic and helper T cells. Latent gammaherpesvirus infection of mice clearly alters autoimmune disease onset and severity reminiscent of the strong association of latent EBV infection in RA patients. As such, these investigative models will serve to identify common mechanisms in which EBV contributes to multiple autoimmune diseases.

Due to EBV infection often taking place years before the onset of arthritis, we posit that latent EBV infection modulates the peripheral immune response in a manner that contributes to the development of RA. We suggest that latently EBV-infected B cells alter, either directly or indirectly, lymphocytes that go on to contribute to disease onset, likely through expanding and activating CD8^+^ T cells and skewing towards a Th1 response. CD11c^+^CD8^+^ DCs may play a role in priming the pathogenic CD8^+^ T cell response, as they have been shown to cross-present antigen(47,48). By acting as a mediator between infected cells and pathogenic T cells, age-associated B cells are likely critical moderators in driving the heightened Th1 immune response to latent viral infection.

Accumulating evidence shows that ABCs are expanded in multiple autoimmune diseases and function pathogenically in mouse models of lupus(25–32). Precisely how ABCs are contributing to pathogenicity is unclear, and ABCs are known to display multiple functional capacities that could contribute to disease. In models of SLE, ABCs have been shown to secrete autoantibodies and compromise germinal centre responses(30). Additionally, ABCs function as excellent antigen-presenting cells(49). In a model of SLE, the ablation of ABCs decreases activated CD4^+^ T cells and IFNγ-CD8^+^ T cells(32). How precisely ABCs are altering the CD8^+^ T cell population, whether they are cross-presenting antigen or impacting the CD8^+^ T cells indirectly, warrants further investigation. Alternatively, ABCs have been shown to secrete regulatory IL10(25,50), suggesting that a portion of the ABC population, or in some individuals or contexts, could function in a protective manner. Further characterization of the phenotype and functional capacities of ABCs in autoimmune patients may help to elucidate their functional role. RA patients who experience a relapse following B cell depletion therapy are more likely to display a reconstitution profile with increased numbers of memory B cells(51). Whether existing therapeutics, such as B cell depletion therapies or other approved drugs for RA, such as Abatacept (CTLA4 Ig), impact the ABC repertoire remains unknown.

Further evaluation of the influence of viral infection in ABC pathogenicity is needed. It is intriguing that ABCs are pathogenic in a genetic model of SLE without the presence of a virus(32), though we observe that latent γHV68 is necessary for the pathogenicity of ABC’s in CIA. This discrepancy indicates that ABCs may be contributing to disease through various mechanisms, or that different contexts can prime ABCs for pathogenicity. The role of ABCs in controlling viral infections is an ongoing topic of study, with multiple papers recently providing compelling evidence that ABCs are critical for an effective anti-influenza response(36) and are required to control of LCMV infection(52), in part through their secretion of antiviral IgG2a. Additionally, the influence of aging on the ABC population and influence on autoimmunity development and progression warrants further study.

In summary, we have developed an in vivo model of EBV’s contribution to RA that recapitulates aspects of human disease. Further, we have examined the role of age-associated B cells and find that they are critical mediators of the viral-enhancement of arthritis.

## Methods

### Mice

Tbet^fl/fl^ × CD19Cre^+/−^ mice were generated by crossing Tbet^fl/fl^ × CD19Cre^+/−^ and Tbet^fl/fl^ × CD19Cre^−/−^. Tbet^fl/fl^ and CD19Cre^+/−^ mice were provided by Dr. Pippa Marrack(32). C57Bl/6 mice were originally purchased from The Jackson Laboratory. All animals were bred and maintained in the animal facility at the University of British Columbia. All animal work was performed per regulations of the Canadian Council for Animal Care (Protocols A17-0105, A17-0184).

### γHV68 and ACRTA-γHV68 infection

γHV68 (WUMS strain, purchased from ATCC) and ACRTA-γHV68 (originally developed by Dr. Ting-Ting Wu, the generous gift of Dr. Marcia A. Blackman)(53), were propagated in Baby Hamster Kidney (BHK, ATCC) cells. Prior to infection, viruses were diluted in Minimum Essential Media (MEM, Gibco) and maintained on ice. Mice (6-to 8-week-old) were infected intraperitoneally (i.p.) with 10^4^ PFU of γHV68 or ACRTA-γHV68 or mock-infected with MEM. No clinical symptoms were observed from viral infections.

### Induction of CIA

On day 35 post-infection, CIA was induced by injection of immunization-grade, chick type II collagen emulsified in complete Freund’s adjuvant (CFA, Chondrex, Inc.) intradermally at the base of the tail, followed by a booster injection of the same emulsion on day 14, as adapted from(54). Each mouse received 0.1 mg chick type II collagen and 0.25 mg CFA at day 0 and 14.

### Evaluation of CIA severity

Clinical signs of CIA were assessed and scored three times per week beginning at the day of CIA induction: 0 = no symptoms, 1 = slight swelling and/or erythema, 2 = pronounced swelling and erythema, 3 = severe swelling, erythema, ankylosis, as adapted from(55). Hind paws were scored individually by a blinded scorer and added for a single score. Day of onset considered two consecutive scoring days of a score of at least 1. The thickness of each hind paw was measured using a digital caliper and the size was expressed as the average thickness of the two paws.

### Tissue harvesting and processing

Mice were anesthetised with isoflurane and euthanized by cardiac puncture. Blood was collected by cardiac puncture into empty sterile tubes and placed on ice until processing, and mice were perfused with 20 ml sterile PBS to allow for synovial fluid harvesting without blood contamination. Inguinal lymph nodes (ILNs) and spleen were extracted and placed into 2 ml sterile PBS and stored temporarily on ice until processing. Synovial fluid was collected by exposing the knee and ankle joints, removing the patellar ligament, and flushing each flexed ankle and knee joint with sterile RNase/DNase-free PBS (Invitrogen) using an 18-gauge needle, adapted from(56,57). Using a 70 μm cell strainer and a 3 ml syringe insert, spleens and ILNs were each mashed through the cell strainer mesh and a single-cell suspension was prepared for each sample. Splenocytes were incubated in ACK lysing buffer for 10 minutes on ice to lyse red blood cells and remaining cells were kept on ice until further use.

### Flow cytometry analysis of cell-type specific surface antigens and intracellular cytokines

To evaluate cytokine production by various cell types, 4 million isolated splenocytes or inguinal lymph node cells were stimulated ex vivo for 3 hours at 5% CO2 at 37°C in Minimum Essential Media (Gibco) containing 10% fetal bovine serum (FBS, Sigma-Aldrich), 1 μl/ml GolgiPlug (BD Biosciences), 10 ng/ml PMA (Sigma-Aldrich) and 500 ng/ml ionomycin (Thermo Fisher). Stimulated cells were then washed prior to staining. For each spleen and ILN sample, 4 million cells were stained in 2 wells, with 2 million cells per well. All collected synovial fluid cells were resuspended in FACS buffer and stained in a single well. Before staining, samples were incubated at 4°C covered from light for 30 minutes with 2 ul/ml Fixable Viability Dye eFluor506 (Thermo Fisher) while in FACS buffer (PBS with 2% newborn calf serum, Sigma-Aldrich). Cells were then incubated with a rat anti-mouse CD16/32 (Fc block) (BD Biosciences) antibody for 10 minutes. Fluorochrome labeled antibodies against cell surface antigens were then added to the cells for 30 minutes covered from light at 4°C. After washing, cells were suspended in Fix/Perm buffer (Thermo Fisher) for 30 minutes-12 hours covered from light at 4°C, washed twice with Perm buffer, and incubated 40 minutes with antibodies for intracellular antigens in Perm buffer. Cells were then washed and resuspended in FACS buffer with 2 mM EDTA. Cells were stained with anti-mouse CD45 (Clone 30-F11, Thermo Fisher Scientific), CD3 (Clone eBio500A2, Thermo Fisher Scientific), CD19 (Clone eBio1D3, Thermo Fisher Scientific), CD4 (Clone RM4-5, Thermo Fisher Scientific), CD8 (Clone 53-6.7, Thermo Fisher Scientific), FoxP3 (Clone FJK-16S, Thermo Fisher Scientific), IFNγ (Clone XMG1.2, Thermo Fisher Scientific), IL17 (Clone TC11-18H10.1, Thermo Fisher Scientific), IL10 (Clone JES5-16E3, Thermo Fisher Scientific), CD11c (Clone 418, Thermo Fisher Scientific), Tbet (Clone eBio4B10, Thermo Fisher Scientific), CD11b (Clone M1/70, Thermo Fisher Scientific), IgD (Clone 11-26c, Thermo Fisher Scientific), CTLA4 (Clone UC10-4B9, Thermo Fisher Scientific), PDL1 (Clone MIH5, Thermo Fisher Scientific), PD1 (Clone J43, Thermo Fisher Scientific), IgM (Clone RMM-1, BioLegend), MHCII (Clone M5/114.15.2, BioLegend), CD27 (Clone LG.3A10, BioLegend), CD20 (Clone SA275A11, BioLegend), TNFα (Clone MP6-XT22, BioLegend), CD95 (Clone SA367H8, BioLegend), and IDO (Clone mIDO-48, Thermo Fisher Scientific). The entirety of each sample was collected on an Attune NxT Flow Cytometer (Thermo Fisher) and analyzed with FlowJo software v10 (FlowJo LLC). Full-minus-one (FMO) controls used for gating.

### IL17 and IFNγ qPCR

RNA was extracted from synovial fluid and inguinal lymph nodes with a Qiagen AllPrep DNA/RNA Micro kit. cDNA was synthesized using Applied Biosystems High-Capacity cDNA Reverse Transcription Kit (Thermo Fisher). qPCR was performed using iQTM SYBR® Green supermix (Bio-Rad) on the Bio-Rad CFX96 Touch™ Real Time PCR Detection system. Primer sets from Integrated DNA Technologies were IL17a 5’-GCT CCA GAA GGC CCT CAG-3’ (forward) and 5’-AGC TTT CCC TCC GCA TTG-3’ (reverse) and IFNγ 5’-ACT GGC AAA AGG ATG GTG AC-3’ (forward) and 5’-TGA GCT CAT TGA ATG CTT GG-3’ (reverse). Normalized to the ribosomal housekeeping gene 18s 5’-GTAACCCGTTGAACCCCATT-3’ (forward) and 5’-CCATCCAATCGGTAGTAGCG-3’ (reverse) and expression determined relative to control group.

### Statistics

Data and statistical analyses were performed using GraphPad Prism software 8.4.2 (GraphPad Software Inc.). Results are presented as mean ± SEM. Statistical tests, significance (p-value), sample size (n, number of mice per group) and number of experimental replicates are stated in the figure legends. Statistical analyses included: two-way ANOVA with Geisser-Greenhouse’s correction, unpaired t-test with Welch’s correction if standard deviations were significantly different, or one-way ANOVA. P-values indicated by asterisks as follows: ****p<0.0001, *** p<0.001, ** p<0.01, * p<0.05.

### Study approval

All work was approved by the Animal Care Committee (ACC) of the University of British Columbia (Protocols A17-0105, A17-0184).

## Author contributions

Experiments were conceived of and designed by ICM and MSH. Experiments conducted by ICM and ZJM. Data analyzed by ICM. IS contributed materials and analysis tools, maintained the mouse colony, and performed genotyping. Paper written by ICM, KLB, and MSH.

## Acknowledgements

We are grateful to Dr. Philippa Marrack for providing Tbet^fl/fl^ and CD19Cre^+/−^ mice and to Jessica R. Allanach for advice and help preparing the manuscript. This research was supported in part by the MSSC, CIHR, UBC, and JDRF.

## Supplementary figures

**Figure 1 – figure supplement 1:**
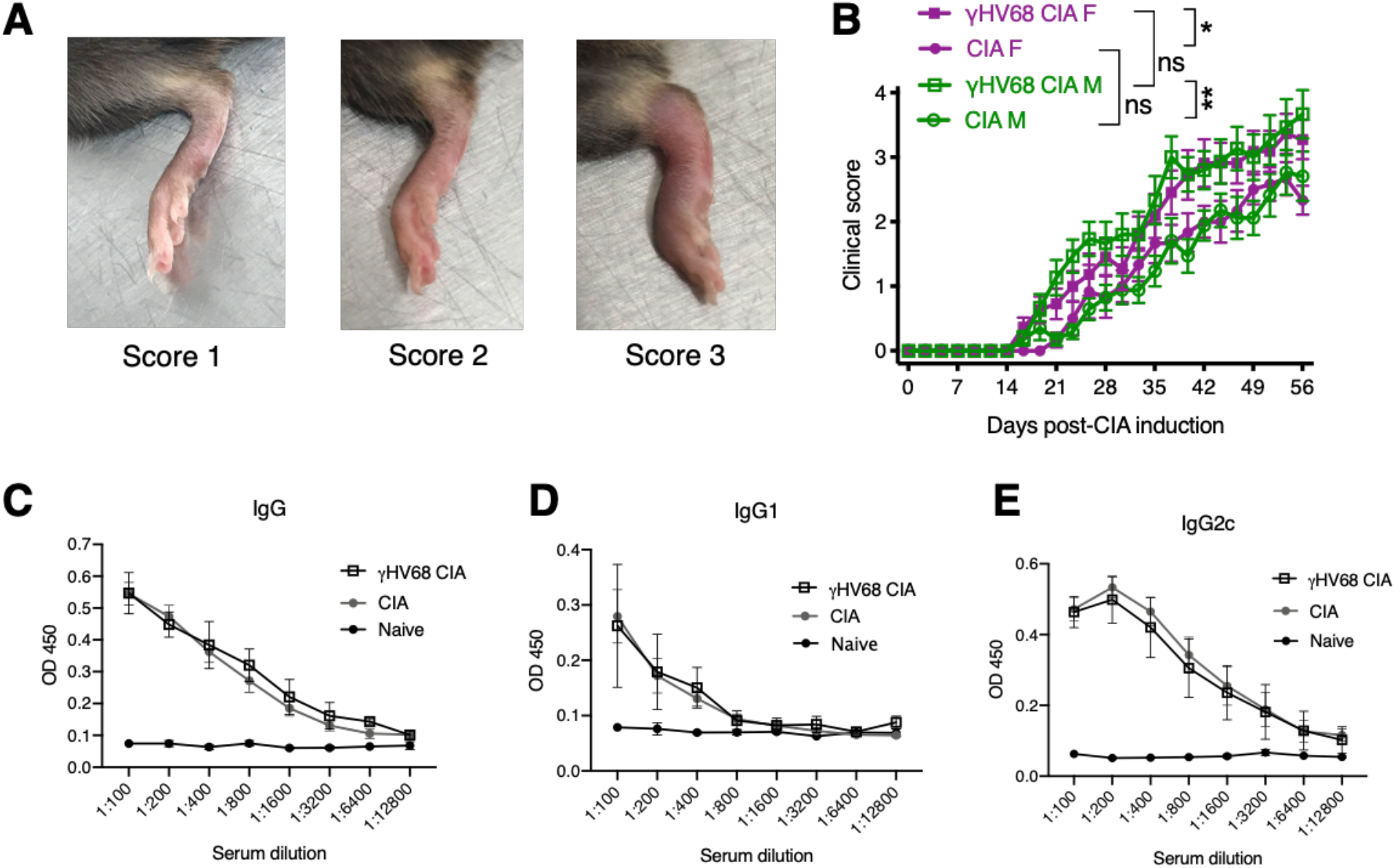
CIA paw scores and autoantibody titers. (**A**) Representative photographs of paws with a corresponding CIA clinical score of 1, 2, and 3. (**B**) Clinical scores (y-axis) over the course of CIA (x-axis, days post induction) in mice (n=11-17 mice per group, same data as in Figure 1A) without (CIA) and with (γHV68-CIA) latent γHV68 infection, separated by sex. Data presented as mean ± SEM and analyzed by two-way ANOVA. ns = not significant. (**C-E**) Optical density (O.D., y-axis) reflecting titers (x-axis; dilution of serum) of anti-type II collagen antibodies separated by (**C**) total IgG, (**D**) IgG1, and (**E**) IgG2c in naïve (black circles), CIA (grey circles), and γHV68-CIA (open squares) mice, n=3-4 mice per group. Data presented as mean ± SEM.

**Figure 3 – figure supplement 1.**
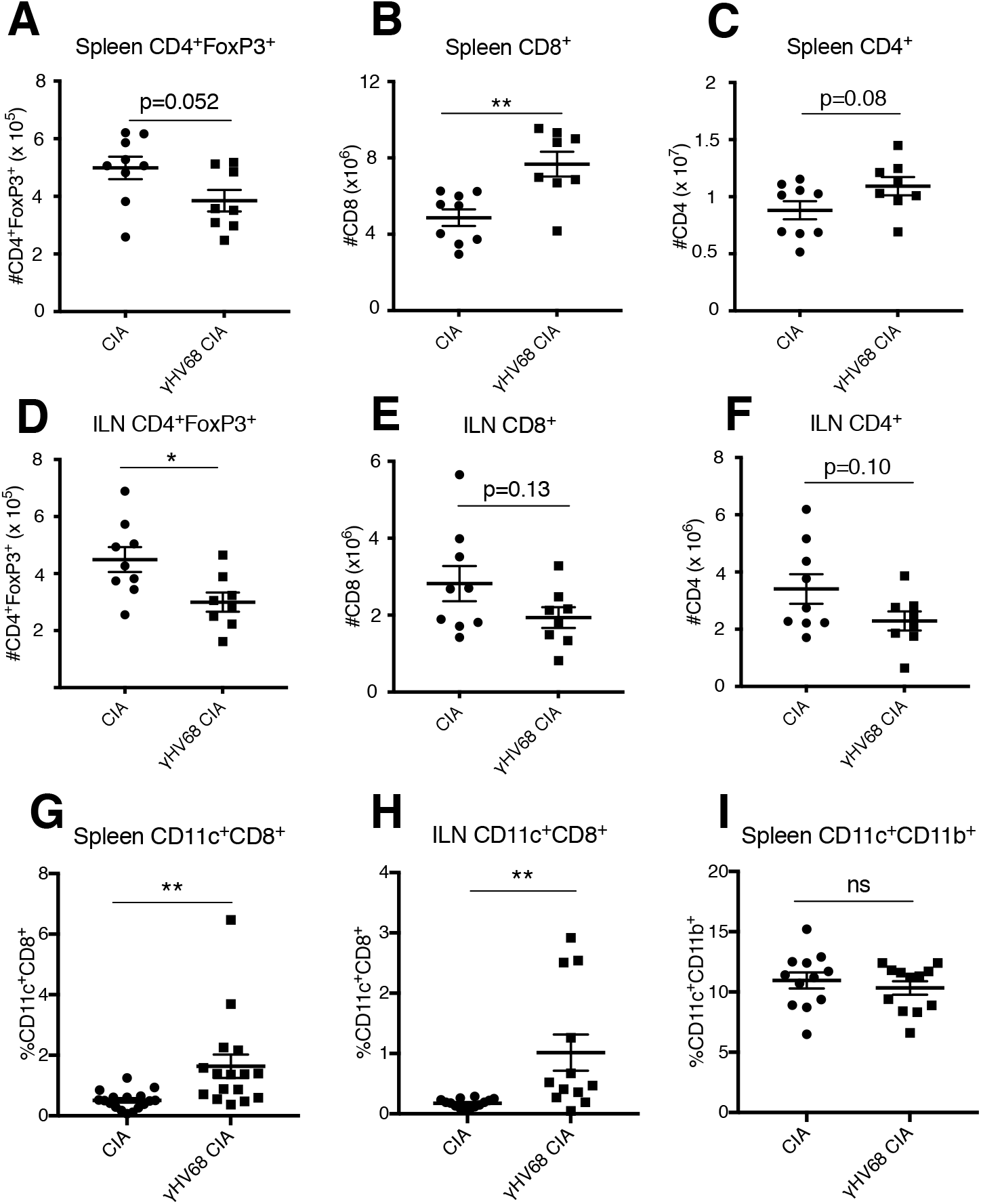
Total numbers of spleen and inguinal lymph node T cell populations at day 56 post-CIA induction. (A-I) Total numbers of T cell subsets (y-axis) in the spleen and inguinal lymph nodes (ILN), as determined by flow cytometry in uninfected mice with CIA (filled circles) and γHV68-CIA mice (filled squares). (**A**) Total number of CD4^+^FoxP3^+^ cells in spleen, (**B**) Total number of CD3^+^CD8^+^ cells in spleen, (**C**) Total number of CD3^+^CD4^+^ cells in spleen. (**D**) Total number of CD4^+^FoxP3^+^ cells in ILN, (**E**) Total number of CD3^+^CD8^+^ cells in ILN, **(F)** Total number of CD3^+^CD4^+^ cells in ILN. Proportion of CD11c^+^CD8^+^ cells of CD45^+^CD3^−^in the spleen and (**H**) ILN. (**I**) Proportion of CD11c^+^CD11b^+^ of CD45^+^CD19^−^CD3^−^in the spleen. (**A-F**) n=8-9 mice per group, 1 experiment, (**G**) n= 16-17 mice per group, 3 experiments (**H**) n=12-14 mice per group, 2 experiments, (**I**) n=12 mice per group, 2 experiments. (**A-I**) Each data point represents an individual mouse. Data presented as mean ± SEM, analyzed by t-test, ** p<0.01, * p<0.05, ns = not significant.

**Figure 5 – figure supplement 1.**
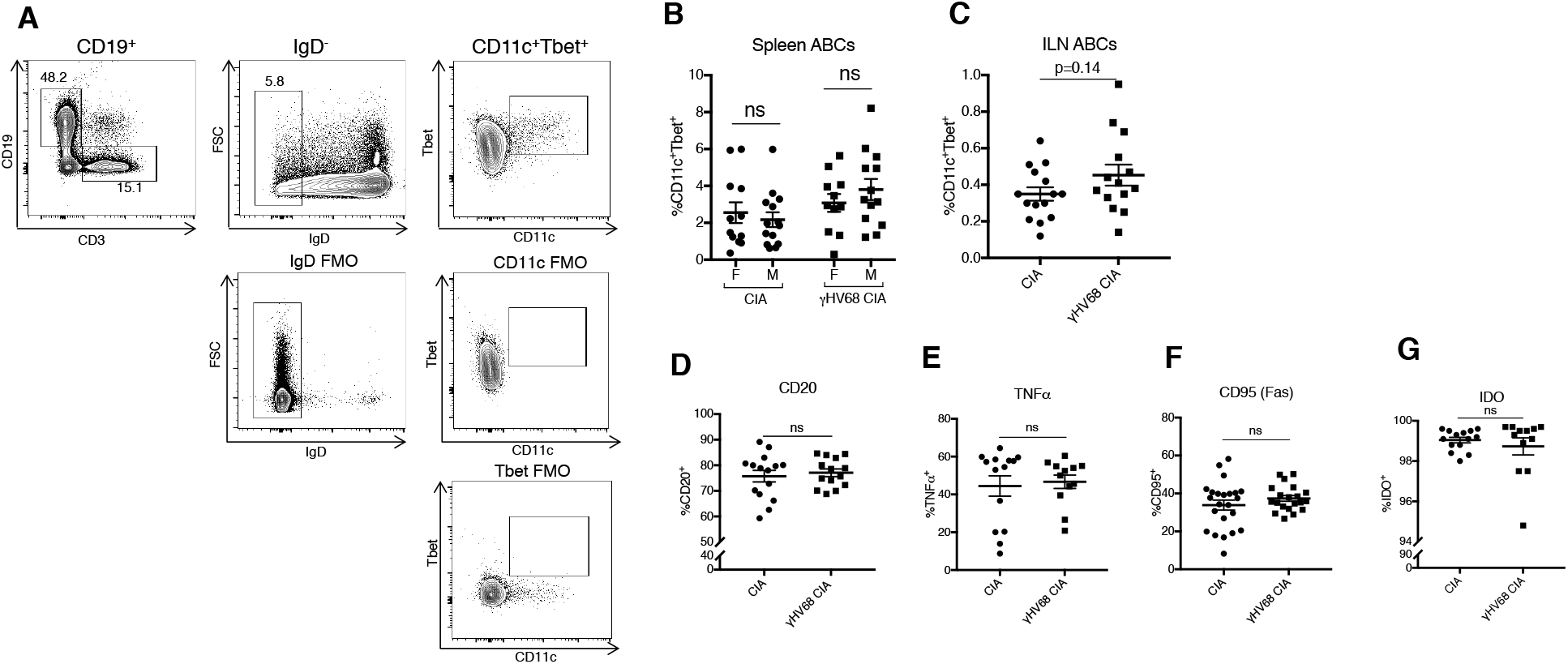
ABCs in the spleen analyzed by flow cytometry at 56 days post-CIA induction. (**A**) Representative gating strategy for ABCs in the spleen using full-minus one (FMO) controls. (**B**) Proportion of ABCs (CD11c^+^Tbet^+^) of mature B cells (CD19^+^IgD^−^) in the spleen of female (F) and male (M) mice in uninfected mice with CIA (filled circles) and γHV68-CIA mice (filled squares). Same results as figure 5A, separated by sex. (**C**) Proportion of ABCs (CD19^+^CD11c^+^) of mature B cells (CD19^+^IgD^−^) in the inguinal lymph nodes (ILNs). (**D-F**) Proportion of ABCs in the spleen expressing (**D**) CD20, (**E**) TNFα, (**F**) CD95, and (**G**) IDO, determined by flow cytometry. Previously gated on ABCs (CD19^+^CD11c^+^Tbet^+^). (**B**) n=11-14 mice per group, 3 experiments, (**C**) n=14-15 mice per group, 2 experiments, (**D**) n=14-15 mice per group, 2 experiments, (**E, G**) n=12-14 mice per group, 2 experiments, (**F**) n=20-23 mice per group, 2 experiments. (**B-G**) Each data point represents an individual mouse. Data presented as mean ± SEM. Analyzed by t-test ****p<0.0001, *** p<0.001, ** p<0.01, * p<0.05, ns = not significant.

## Supplementary methods

### Anti-type II collagen antibody ELISA

The sera were isolated by centrifugation 2000 × g for 10 min, aliquoted, and stored for up to 14 months at −80°C prior to running the ELISA. Anti-type II collagen antibodies were quantified by standard indirect ELISA. Briefly, ELISA plates (NUNC, Thermo Fisher) were coated with 5 μg/ml ELISA-grade type II collagen (Chondrex, Inc.) overnight at 4°C, washed 4x with wash buffer (PBS, 0.05% Tween-20), blocked with 5% newborn calf serum (NBCS, Sigma-Aldrich) for 1 hour at 37°C, incubated with serial dilutions (1:100 to 1:12800) of test sera diluted in blocking buffer for 2 hours at 37°C, and washed 4x wash buffer. Bound (anti-collagen II) antibody was incubated with HRP-conjugated goat anti-mouse IgG (Thermo Fisher), rat anti-mouse IgG1 (BD Biosciences), or goat anti-mouse IgG2c (Thermo Fisher), all diluted 1:500 in blocking buffer, for 1 hour at 37°C, washed 4x with wash buffer, and detected by TMB substrate (BD Biosciences). Absorbance was read at 450 nm on a VarioSkan Plate Reader (Thermo Fisher).

